# crispAIPE: Probabilistic Modelling of Prime Editing Variant Correction Efficiency

**DOI:** 10.64898/2025.12.07.692852

**Authors:** Furkan Özden, Peiheng Lu, Peter Minary

**Affiliations:** Department of Computer Science, University of Oxford, Parks Rd, Oxford OX1 3QD, United Kingdom

**Keywords:** Genome Editing, Prime Editing, Uncertainty Quantification, Conformal Prediction, Deep Learning, Transformers, CRISPR

## Abstract

Prime editing installs precise substitutions, insertions, and deletions without double-strand breaks or donor templates, but the efficiency of an individual pegRNA design is hard to predict in advance, and existing tools give only point estimates, leaving designers unable to judge which predictions to trust. We present crispAIPE, a transformer-based probabilistic framework that quantifies per pegRNA–target pair uncertainty by modelling the three competing prime-editing outcomes on the 2-simplex with a Dirichlet likelihood and pairing the posterior with split-conformal highest-density regions calibrated on a held-out fold, giving finite-sample coverage guarantees on the simplex without requiring the Dirichlet model itself to be well-calibrated. Trained on 92,423 PRIDICT Library-1 pegRNAs under mutation-level target-disjoint splitting, crispAIPE attains Spearman *ρ* = 0.835, 0.843, and 0.693 on the edited, unedited, and indel fractions (Pearson *r* = 0.845, 0.855, 0.659), and its conformal regions match nominal coverage where region-construction baselines do not. pegRNA architecture and edit context, in particular GC content of the mutated reverse transcription template (RTT) and edit size for deletions, are associated with prediction uncertainty. Reusing the Library-1 calibration quantile unchanged on PRIDICT Library-2, the conformal region area generalises as an actionable filter for cross-cell-type design, identifying the pegRNAs whose Library-1 predictions transfer best while small-sample head fine-tuning further improves accuracy. Tool and trained models: https://github.com/furkanozdenn/pe-uncert.

## 1 Introduction

Prime editing is a CRISPR-derived genome-editing modality that installs precise, user-specified DNA changes (single-base substitutions, small insertions, and small deletions) directly into a chosen locus without creating a double-strand break and without a separately delivered donor template [5]. Mechanistically, a prime editor protein (a partially catalytically dead Cas9 nickase fused to an engineered reverse transcriptase) is loaded with a *prime editing guide RNA* (pegRNA), an extended single-guide RNA whose 5^*′*^ *spacer* (∼ 20 nt) directs the editor to the genomic *protospacer*, and whose 3^*′*^ extension contains a *primer binding site* (PBS, ∼ 8–17 nt) and a *reverse transcription template* (RTT, ∼ 10–20 nt) that together encode the new sequence. The portion of the RTT downstream of the intended edit is the *RTT overhang*, which provides the homology needed to commit the new flap to the genome; varying RTT overhang length, PBS length, and edit position within the RTT defines the principal pegRNA design space. Editor evolution has progressed from PE1 (the original Cas9 nickase + wild-type M-MLV reverse transcriptase [5]) and PE2 (engineered M-MLV reverse transcriptase [5]) through PE3/PE3b (PE2 + a nicking guide RNA on the unedited strand) [5], PE4/PE5 (transient mismatch-repair inhibition via dominant-negative MLH1) [8], PE6 (phage-evolved compact reverse transcriptases) [11], and PE7 (an editor–La fusion that stabilises the pegRNA 3^*′*^ end) [39]. PRIDICT Library-1, the screen used in this work, was generated with PE2 in HEK293T cells and remains the de facto benchmark for prime-editing efficiency prediction [28].

Prime editing has shown therapeutic potential for correcting disease-causing mutations [14, 35], with preclinical demonstrations in sickle cell disease [13, 25] and cystic fibrosis [37] reaching clinically relevant efficiency, and a first in-human trial for p47^phox^-deficient chronic granulomatous disease showing DHR-positive neutrophils above the clinical-benefit threshold within thirty days. The platform has evolved rapidly: PE4/PE5 [8], engineered pegRNAs [30], PE6 [11], and PE7 [39], and now supports large deletions [9], extended [4] and gene-sized insertions [32], and multiplexed editing [16]. Yet success depends critically on pegRNA design [7, 12]: efficiency varies across target sites and genomic contexts [23] through a complex multi-step mechanism influenced by mismatch repair, chromatin accessibility, and cell-cycle phase [41]. Computational predictors have grown from DeepPE [19] and PRIDICT [28] to DeepPrime [40], PRIDICT2.0 [29], and OPED [27], with specialised tools for insertion efficiency [21] and chromatin integration [26].

A critical limitation of existing tools is their reliance on point predictions without uncertainty quantification, which precludes assessing prediction reliability when selecting pegRNAs for validation, especially for clinical use, where distinguishing confident from uncertain predictions is essential. Uncertainty quantification is standard in high-stakes domains such as medical diagnosis and autonomous systems [6, 18], reduces error under distribution shift in biomedical AI [10, 33], and has proven valuable for CRISPR off-target assessment [31], yet no tool provides uncertainty estimates for prime-editing efficiency. Here we present crispAIPE (CRISPR-AI Prime Editing), the first uncertainty-aware framework for prime-editing efficiency prediction: built on PRIDICT datasets [28], it uses probabilistic neural networks to estimate posterior predictive uncertainty, providing full outcome distributions rather than point estimates to support reliability-aware pegRNA prioritization.

## 2 Materials and Methods

### 2.1 Datasets

We trained and evaluated crispAIPE on PRIDICT Library-1 [28], a high-throughput prime editing screen targeting 13,349 pathogenic ClinVar mutations [22], encompassing single base replacements and insertions (up to 24 bp) and deletions (up to 13 bp). PRIDICT Library-1 is a lentiviral self-targeting library in which each pegRNA is paired in cis with its own target site [28]; after seven days of editing with PE2 in HEK293T cells, genomic DNA is harvested and the integrated cassette is read out by amplicon high-throughput sequencing. For every pegRNA–target pair, sequencing reads are classified into one of three mutually exclusive categories: carrying the intended edit (*edited*), remaining unmodified (*unedited*), or carrying any unintended modification (*indel*). The unintended-edit category aggregates *all* reads that deviate from both the wild-type and the intended-edit sequence, i.e. unintended insertions, deletions, and substitutions outside the intended position; following PRIDICT [28] we refer to this fraction as the *indel* fraction for brevity, although it is not restricted to insertions and deletions. The per pegRNA–target pair outcome is the read-fraction triplet (*y*_*e*_, *y*_*u*_, *y*_*i*_) summing to one, averaged across three biological replicates (Spearman *R >* 0.98, Pearson *r >* 0.99 across replicates). After quality control filtering of an initial 119,701 pegRNA–target pairs (excluding pegRNAs with fewer than 100 reads per replicate and pegRNAs whose spacer did not match the target), 92,423 high-quality pairs remained (57,920 substitutions, 28,420 insertions, 6,083 deletions), with a median edited fraction of 46% and a median indel fraction of 7.4% [28]. crispAIPE is trained on these three-outcome proportions directly via a Dirichlet likelihood (Section 2.2), so the prediction target is the joint outcome composition rather than a single scalar efficiency. Cell-line transfer analyses used PRIDICT Library-2 conditions reported by Mathis et al. For the regression variant (crispAIPE-reg), we used the HEK293T PE2 subset of DeepPrime [40], comprising 259,910 training and 28,883 test pegRNAs.

*Target-disjoint data splitting*. The PRIDICT Library-1 dataset contains multiple pegRNAs per target mutation differing primarily in RTT overhang length while sharing the same target site and sequence context. Random splitting of such related pegRNAs across partitions constitutes data leakage and can inflate performance metrics. We therefore used the original PRIDICT mutation identifier as the grouping variable and assigned all pegRNAs targeting the same mutation to the same partition. Groups were stratified by edited-fraction quartile before splitting to preserve the outcome distribution across partitions. This yielded 64,751 training, 13,828 validation, and 13,844 test samples with zero mutation-level target overlap between partitions.

### 2.2 Modelling of crispAIPE

#### 2.2.1 Problem formulation

Let **x**_init_, **x**_mut_ ∈ {0, 1}^*m×L*^ be the one-hot encoded wild-type (initial) and mutated (edited) target sequences, each of fixed length *L* = 99, where *m* = 5 is the alphabet size: the four nucleotides {A, C, G, T} together with a gap symbol used to pad insertions and deletions so that the initial and mutated sequences share the common length *L*. Let **r** ∈ {0, 1}^4*×L*^ denote binary positional masks for the protospacer, PBS, RTT-initial, and RTT-mutated regions (one channel each). We write the complete model input as the triple **x** = (**x**_init_, **x**_mut_, **r**) ∈ *X*, where the input space is *X* = {0, 1}^*m×L*^ × {0, 1}^*m×L*^ × {0, 1}^4*×L*^.

From **x** a unified representation **u** ∈ {0, 1}^11*×L*^ is formed by row-wise concatenation

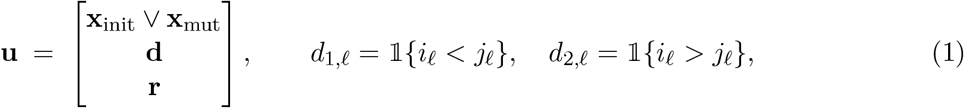

where ∨ denotes element-wise logical OR (5 channels) and *i*_*ℓ*_, *j*_*ℓ*_ ∈ {1, …, 5} are the non-zero indices of the *ℓ*-th columns of **x**_init_ and **x**_mut_ under a fixed alphabet ordering, so that **d** ∈ {0, 1}^2*×L*^ encodes mismatch directionality (2 channels) and the masks **r** contribute the remaining 4 channels.

The compositional editing outcome **y** = (*y*_*e*_, *y*_*u*_, *y*_*i*_) lies on the 2-simplex 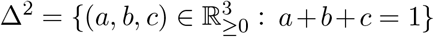 and gives the proportions of edited, unedited, and indel-containing reads respectively.

We model the outcome distribution using a Dirichlet, which naturally captures the competitive relationship between the three editing outcomes and varying experimental precision through its concentration parameters. We learn a parametric function 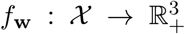 mapping inputs to concentration parameters ***α*** = (*α*_*e*_, *α*_*u*_, *α*_*i*_):

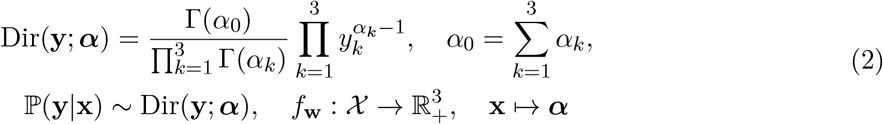

The expected efficiency is 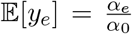 and variance 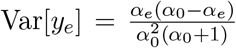, enabling both point estimates and posterior uncertainty diagnostics over editing outcomes.

#### 2.2.2 Encoding and Tokenization

We employ a hybrid encoding strategy combining 5-bit nucleotide encoding, 2-bit edit directionality channels, and 4-bit functional region annotations (protospacer, PBS, RTT-initial, RTT-mutated) to create an 11 × *L* unified representation. This multi-channel representation captures sequence content, edit positions and directions, and pegRNA functional elements through element-wise OR operations and binary masks (Supplementary Note 2). In crispAIPE, the wild-type nucleotide sequence is also converted to 5-symbol token indices and passed through a learnable nucleotide embedding.

#### 2.2.3 Architecture and Training of crispAIPE

crispAIPE is an end-to-end probabilistic neural network combining transformer encoders with convolutional layers to predict prime editing outcomes via Dirichlet distributions. Nucleotide tokens are mapped to dense embeddings (*d*_embed_ = 64) with sinusoidal positional encoding. Two stacked transformer encoder layers with four attention heads (*d*_*ff*_ = 32, *p*_*drop*_ = 0.1) produce **h**_*tr*_ ∈ ℝ^*L×*64^, which is concatenated with the transposed unified representation **u**^*T*^ ∈ ℝ^*L×*11^ to form **h**_combined_ ∈ ℝ^*L×*75^. The combined representation is passed through a one-dimensional convolutional block with 64 output channels, ReLU activation, batch normalization, and global max pooling. A final multilayer perceptron maps the pooled representation to three Dirichlet concentration parameters: 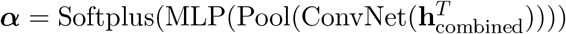, where Softplus ensures *α*_*i*_ *>* 0.

The model minimises the negative log-likelihood of the Dirichlet distribution:

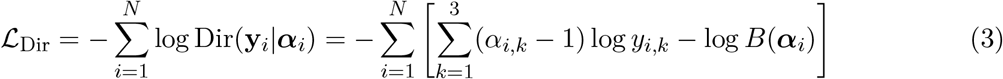

where *B*(***α***) is the multivariate Beta function. We used Adam [20] with learning rate 6 × 10^−4^, cosine decay with 10 warm-up epochs, batch size 128, a maximum of 100 epochs, and early stopping with patience of 8 epochs in PyTorch [34]. Target-disjoint analyses use the hybrid transformer– CNN Dirichlet model as the primary model. Architecture and outcome-parameterisation ablations were trained on the same mutation-level target-disjoint split (Supplementary Tables S1–S2).

#### 2.2.4 Conformal Highest-Density Region Construction

To quantify prediction uncertainty, crispAIPE constructs prediction regions on the 2-simplex Δ^2^ representing the three editing outcomes (edited, unedited, indel) using split conformal prediction [3, 24, 38] with a Dirichlet highest-density region (HDR) geometry [2, 17]. The region geometry follows Hyndman’s density-quantile HDR: for a test input **x**, the region is the super-level set 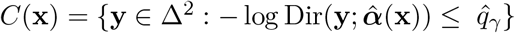. The threshold 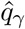 is calibrated on a held-out fold rather than read off the model’s own posterior. Concretely, for each pegRNA–target pair 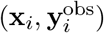 in the validation calibration fold (*n*_cal_ = 13 828) we compute the nonconformity score 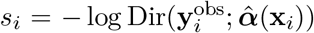 and set 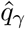 to the ⌈ (*n*_cal_ + 1)(1 − *γ*) ⌉*/n*_cal_ empirical quantile of {*s*_*i*_} . Under exchangeability of calibration and test points, this guarantees P(**y**^obs^ ∈ *C*(**x**)) ≥ 1 − *γ* in finite samples [24, 38], irrespective of whether 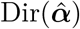 is well-calibrated. The region is a level set of the Dirichlet density and therefore stays on the simplex by construction. Supplementary Note 1 details the construction. The resulting calibration comparison against a Bayesian HDR on 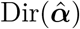 and a PCA-ellipse baseline is shown in the main text (Fig. 1C, D), with the full region-construction ablation in Supplementary Fig. S4.

**Figure 1.**
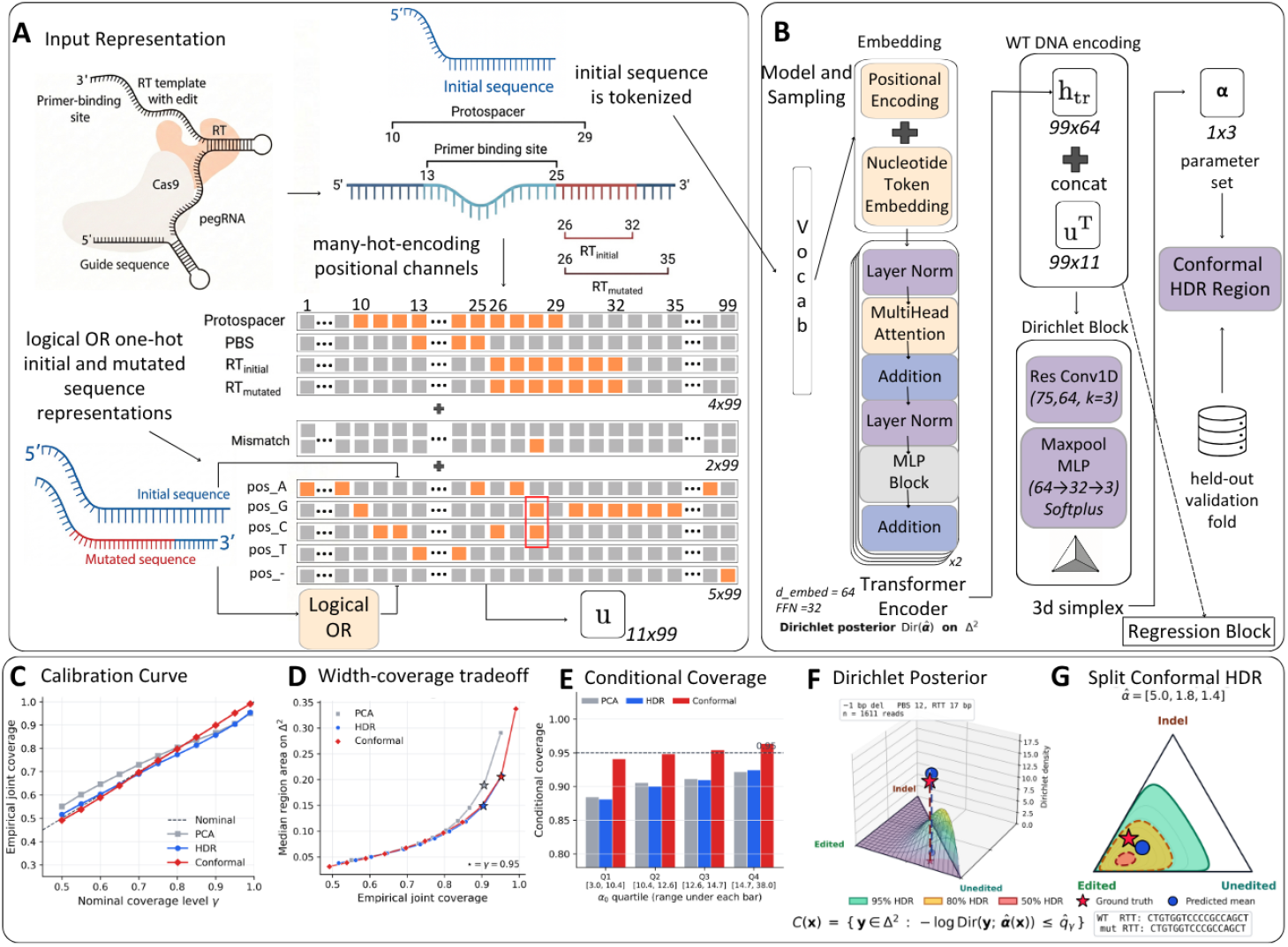
crispAIPE: a transformer-based framework for uncertainty-aware prime-editing efficiency prediction. **(A)** Input-representation pipeline: wild-type and mutated sequences for a pegRNA–target pair are encoded into nucleotide tokens and an 11 × *L* unified representation (*L* = 99) capturing guide, primer binding site (PBS), reverse-transcription (RT) template positions, and mismatch directions. **(B)** Model and sampling architecture: nucleotide embeddings pass through transformer encoder layers, are concatenated with the unified representation, and are mapped by convolutional and MLP layers to Dirichlet concentration parameters ***α*** = [*α*_*e*_, *α*_*u*_, *α*_*i*_]; the predicted Dirichlet feeds the split-conformal HDR constructor whose threshold 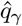 is calibrated once on the held-out validation fold. The separate crispAIPE-reg regression block (MSE-trained, single edited-efficiency output) is indicated and can replace the Dirichlet head for point prediction. **(C)** Calibration curve: empirical versus nominal joint coverage for the split-conformal HDR against PCA-ellipse and Bayesian-HDR baselines. **(D)** Width–coverage trade-off: median region area on Δ^2^ versus empirical coverage, with stars marking the *γ* = 0.95 operating point. **(E)** Conditional coverage at *γ* = 0.95 across Dirichlet-concentration *α*_0_ quartiles. **(F)** Dirichlet posterior density for a representative high-efficiency pe-gRNA (a 1 bp deletion, RTT 17 bp, *n* = 1,611 reads, observed edited fraction 0.66, 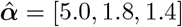), with ground truth (red star) and predicted mean (blue circle). **(G)** Equilateral-ternary view of Δ^2^ for the same pegRNA with nested 50% (red), 80% (amber), and 95% (green) split-conformal HDR regions, each defined as 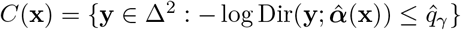.

## 3 Results

### 3.1 Overview of crispAIPE

We developed crispAIPE, a transformer-based architecture for uncertainty-aware prime-editing prediction (Figure 1) that models editing outcomes as a distribution over the edited, unedited, and indel fractions with a Dirichlet framework.

Figure 1 summarises the workflow: wild-type and mutated sequences are encoded into nucleotide tokens and an 11 × *L* unified representation (Fig. 1A); the model returns Dirichlet concentration parameters 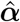 (Fig. 1B), and a split-conformal HDR constructor turns these into per pegRNA–target pair prediction regions on the simplex with finite-sample marginal coverage, using a single calibration quantile 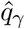 fitted on the held-out validation fold. Supplementary Fig. S2 shows representative posteriors and conformal regions across editing efficiencies.

### 3.2 crispAIPE enables uncertainty quantification with superior predictive power in prime editing outcomes

On the target-disjoint Library-1 test set (*n* = 13,844), crispAIPE predicts all three outcome fractions accurately (Fig. 2A–C): Spearman *ρ* = 0.835 (edited), 0.843 (unedited), and 0.693 (indel; limited by the lower signal and smaller dynamic range of indel outcomes), for an overall *ρ* = 0.851 (*r* = 0.863). Against published point-prediction benchmarks (Fig. 2D), crispAIPE is competitive with PRIDICT and OPED on the edited (“intended”) and indel (“unintended”) fractions; the comparison is only approximate because PRIDICT uses a locus-grouped split comparable to ours, whereas OPED’s reported values come from a random 80/20 split of Library-1 that does not exclude same-target pegRNAs across train and test (asterisked in Fig. 2D). Part of this gap is expected: PRIDICT and OPED augment the sequence with engineered biophysical features (e.g., pegRNA minimum free energy and melting temperatures), whereas crispAIPE uses sequence and edit context only; matching their rank correlation without such hand-crafted features is a strength, and adding them is a natural extension. On the separate edited-efficiency regression task (DeepPrime test data), crispAIPE-reg reaches Spearman *ρ* = 0.815 (*r* = 0.792; Supplementary Fig. S1), above DeepPrime (0.740) and EasyPrime (0.670). Unlike these point predictors, crispAIPE additionally returns posterior concentration parameters for uncertainty diagnostics.

**Figure 2.**
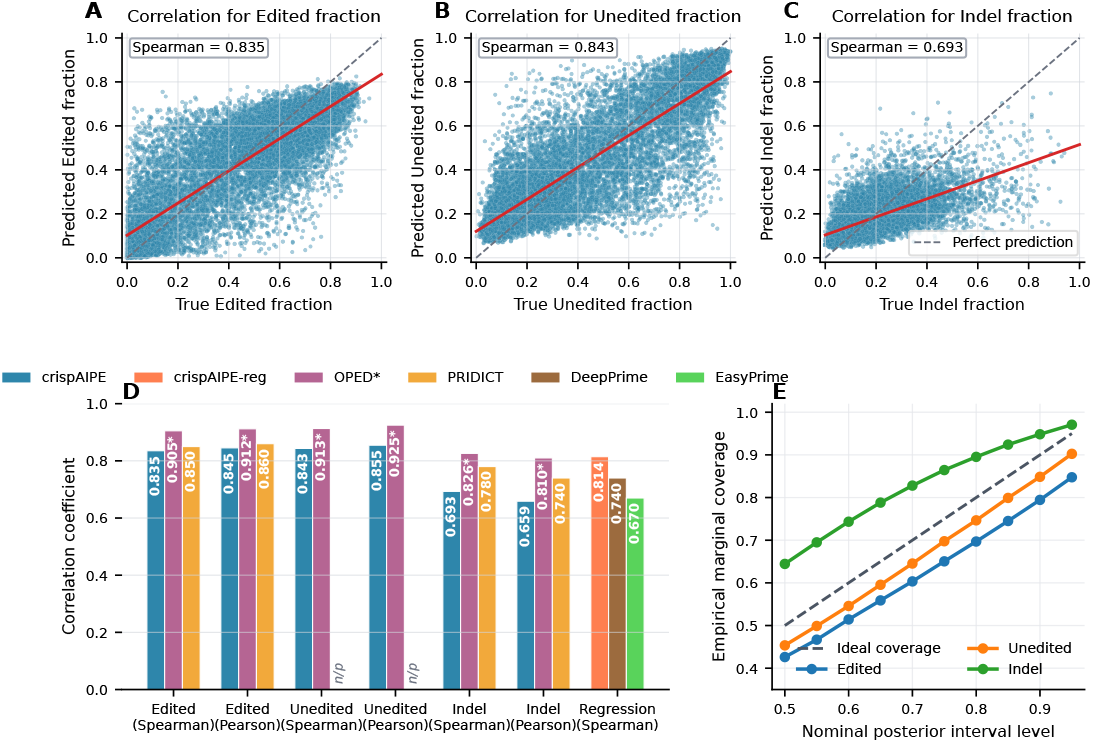
Prediction accuracy, benchmarking, and uncertainty diagnostics for crispAIPE across prime editing outcomes. **(A–C)** Scatter plots showing predicted versus true values for edited, unedited, and indel outcomes on the mutation-level target-disjoint Library-1 test dataset. Each point represents a single pegRNA design; the dashed diagonal indicates perfect prediction and the red line shows the fitted trend. **(D)** Per-outcome Spearman and Pearson correlations for crispAIPE on edited, unedited, and indel fractions, plus the separate edited-efficiency regression Spearman for crispAIPE-reg. PRIDICT’s and OPED’s published “intended” and “unintended” correlations map respectively to the edited and indel rows here; OPED’s unedited comparator is from its Extended Data Fig. 4b (*r* = 0.925, *R* = 0.913), while PRIDICT does not report unedited separately (n/p). PRIDICT is evaluated under locus-grouped splitting comparable to our mutation-level target-disjoint split, whereas OPED’s reported values come from a random 80/20 split of Library-1; OPED’s per-outcome bars carry an asterisk to flag this residual same-target leakage. Regression-task comparators are DeepPrime and EasyPrime. **(E)** Empirical marginal coverage of posterior intervals across nominal interval levels for edited, unedited, and indel outcomes; the dashed line indicates ideal empirical coverage.

Architecture ablations support the hybrid transformer–CNN design (Supplementary Table S1), and distribution-head ablations show a softmax point head is marginally higher in rank correlation (*ρ* = 0.856) than the Dirichlet head (0.851) while a logit-normal is lower (0.829; Supplementary Table S2); we retain the Dirichlet head because it alone yields the concentration parameters and posterior samples needed for the uncertainty diagnostics (Fig. 2E), reported as an empirical coverage check rather than a proof of perfect calibration.

### 3.3 Concentration parameters provide empirical uncertainty diagnostics correlated with prediction accuracy

We assessed crispAIPE’s uncertainty estimates (Figure 3). Across the target-disjoint test set the 95% conformal-HDR region area had median 0.206 (of a maximum 0.5 on the (edited, unedited) projection; Fig. 3A), and empirical joint coverage matched nominal at every level (95.2% at *γ* = 0.95; 49.2/79.7/89.9% at *γ* = 0.50/0.80/0.90, all within 1%; Supplementary Fig. S4). The underlying Dirichlet posterior is marginally miscalibrated, under-covering edited/unedited and over-covering indel (84.8/90.2/97.1% at the 95% Beta-marginal level; Fig. 3B), which the conformal layer absorbs at the joint level. Both quantities track reliability per pegRNA: across the test set (*n* = 13,844), region area is positively correlated with absolute prediction error (Spearman *ρ* = 0.431) and the Dirichlet concentration *α*_0_ negatively so (*ρ* = −0.308): higher-concentration predictions have smaller regions and lower error (Fig. 3C, D), making both empirical reliability indicators for pe-aagRNA prioritization.

**Figure 3.**
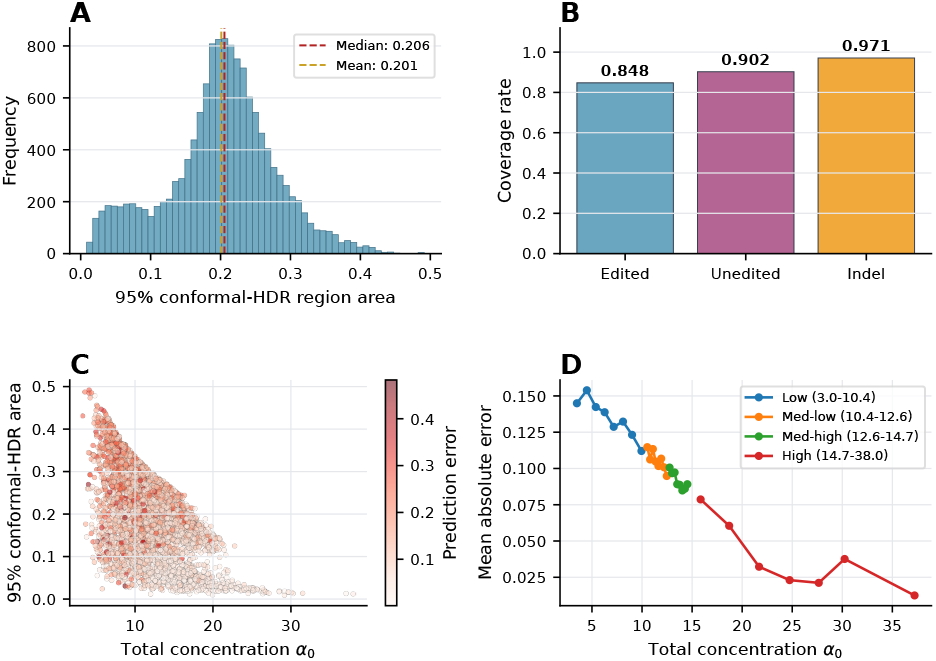
Uncertainty quantification under the split-conformal HDR construction. **(A)** Distribution of 95% conformal-HDR region areas on the (edited, unedited) projection of the simplex across the target-disjoint test set (*n* = 13,844), with median area = 0.206 and mean area = 0.201. **(B)** Empirical marginal coverage rates of the underlying Dirichlet posterior at the 95% level for each outcome (84.8% / 90.2% / 97.1% for edited / unedited / indel); these document the residual marginal miscalibration of the Dirichlet head that the conformal construction absorbs at the joint level. **(C)** Relationship between 95% conformal-HDR area, total Dirichlet concentration *α*_0_, and absolute prediction error. Each point represents a single pegRNA, colored by absolute prediction error. **(D)** Mean absolute error stratified by *α*_0_ quartile, where *α*_0_ = *α*_*e*_ + *α*_*u*_ + *α*_*i*_. Legend labels report the *α*_0_ range of each quartile.

### 3.4 Calibrated outcome regions for pegRNA prioritization

Reliable per pegRNA–target pair uncertainty is what distinguishes prime-editing prediction frameworks that can be acted on from those that cannot. We therefore characterized how the conformal-HDR regions behave across nominal levels and across the heteroskedastic regime indexed by Dirichlet concentration *α*_0_ (Figure 1C–G). The calibration curve (Fig. 1C) shows that the empirical fraction of test pegRNAs whose experimentally observed outcome falls inside the predicted region matches the nominal *γ* across *γ* 0.50, 0.55, …, 0.99 with maximum absolute deviation below 1%, whereas the PCA-ellipse and Bayesian-HDR baselines under- or over-cover by several percentage points at every level. Operationally, selecting pegRNAs whose 95% conformal region lies entirely above a desired edited-fraction threshold gives the experimentalist a frequentist statement: at most 5% of selected designs will fall short. The width–coverage trade-off (Fig. 1D) makes the cost explicit: the Bayesian HDR is sharper at any fixed *γ* but operates strictly below the nominal coverage line, while the conformal HDR is modestly wider in exchange for a real coverage statement. Conditional coverage by *α*_0_ quartile (Fig. 1E) confirms that the guarantee holds across the heteroskedastic regime: empirical coverage at *γ* = 0.95 stays in [0.94, 0.97] in every quartile, whereas the baselines drift below nominal in the low-*α*_0_ stratum. A representative high-efficiency pegRNA (a −1 bp deletion, RTT 17 bp, *n* = 1,611 reads, observed edited fraction 0.66) illustrates the construction at the single-design level: its Dirichlet posterior density (Fig. 1F) is sharp and edited-dominant 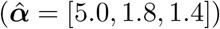, and the nested 50*/*80*/*95% split-conformal regions on the simplex (Fig. 1G) contain the experimentally observed outcome, with the 95% region widening where the point prediction is least certain to provide an actionable flag for further validation. A bootstrap of the calibration quantile (200 resamples of the validation fold) yielded 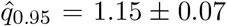 with 95% confidence interval [1.01, 1.27], confirming that calibration is stable at the current calibration size and can be deployed routinely on new prime-editing screens of comparable scale.

## 4 Discussion and Conclusions

Computational prime-editing tools have progressed from DeepPE [19] through PRIDICT [28] (*R* = 0.85) to DeepPrime [40] and PRIDICT2.0 [29], covering diverse edit types and cell lines but providing point predictions without model-specific posterior uncertainty over the edited/unedited/indel composition, which limits per-design reliability assessment. Uncertainty quantification is valuable wherever prediction reliability drives decisions [6, 18], including CRISPR off-target assessment [31] and guide selection [36]; in prime editing the added challenge is that the three competing outcomes are compositional and must sum to unity, a constraint independent regressors do not enforce [1, 15].

Our Dirichlet framework captures this compositional structure with a transformer network paired with split-conformal prediction [3, 24, 38] under a Dirichlet-HDR geometry [2, 17]. Under target-disjoint evaluation the conformal layer attains nominal joint coverage where a Bayesian HDR under-covers and a PCA ellipse leaks off the simplex (Supplementary Fig. S4). The calibrated region area doubles as a feature-level uncertainty map, dominated by mutated-RTT GC content, deletion size, and edit position (Supplementary Fig. S5), and paralleled by transformer attention summaries (Supplementary Fig. S3) that reflect the behaviour of the trained predictor, and as an actionable cross-cell-line transfer filter (Supplementary Fig. S6).

The design space at a single locus reaches 10^3^–10^4^ candidates that cannot all be screened, so beyond ranking by expected efficiency crispAIPE’s per-design Dirichlet posterior and calibrated 95% HDR let an experimentalist rank by upper-confidence edited fraction, flag candidates whose region overlaps low-efficiency outcomes for extra screening before stem- or primary-cell use, and allocate validation effort in proportion to predicted uncertainty; the Library-2 transfer signal (Supplementary Fig. S6) further indicates when published predictions will translate to a new cell line. In practice, crispAIPE helps prioritise which pegRNAs to validate experimentally. Future extensions such as chromatin accessibility, diverse cell-type training data, recalibration, and genomic language-model transfer may further improve deployment.

## Funding & rights retention

This work is funded by Google DeepMind; for Open Access, a CC BY licence applies to any Author Accepted Manuscript from this submission.

## Code & data

https://github.com/furkanozdenn/pe-uncert.

## Author contributions

FO and PM designed the study; PL and FO prepared the datasets; FO implemented the model and experiments; FO and PM wrote the paper.

