## Supplementary Material for "crispAIPE: Probabilistic Modelling of Prime Editing Variant Correction Efficiency"

### Supplementary Figures

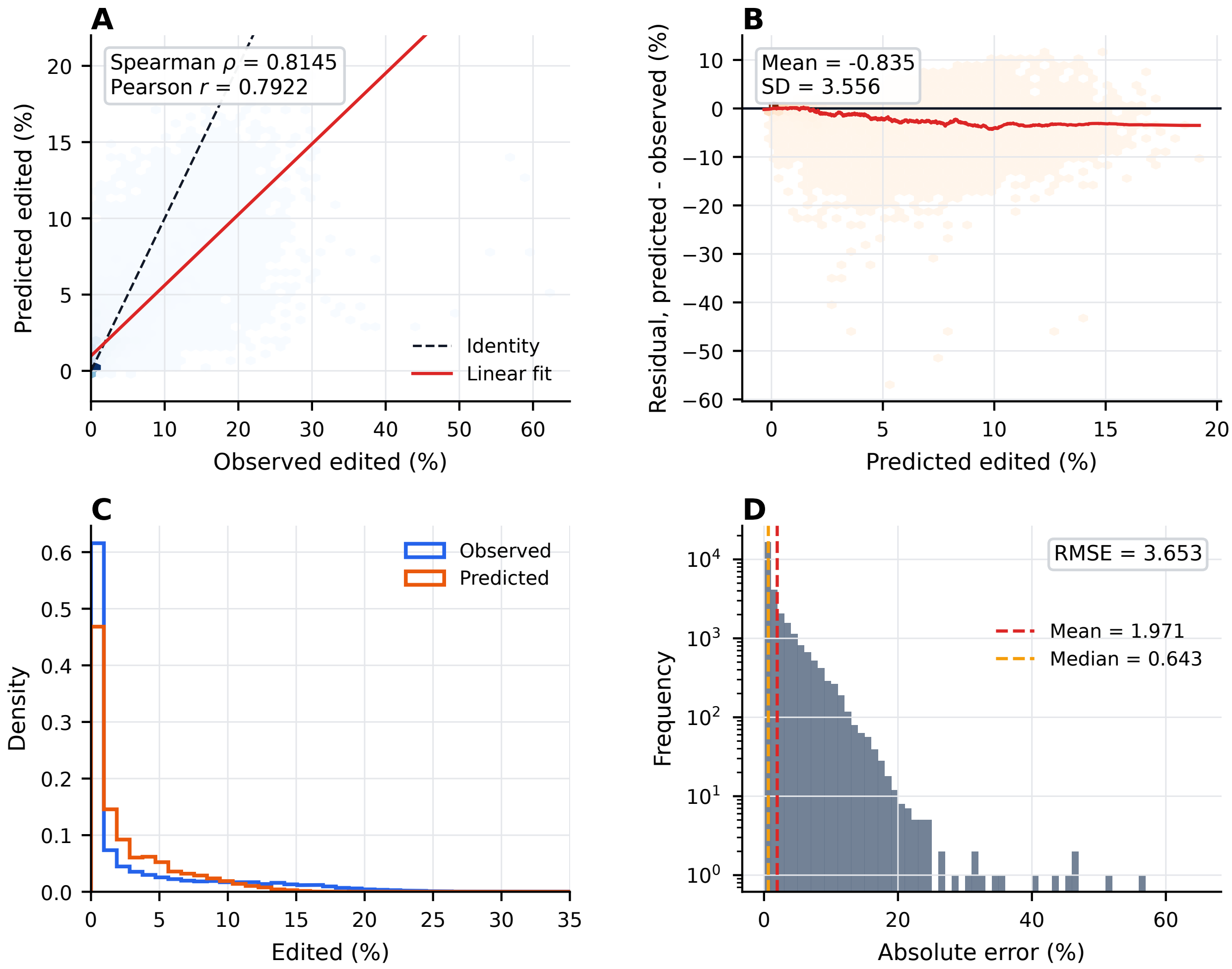


Figure S1: Regression analysis of crispAIPE-reg model performance on the DeepPrime test dataset ($n = 28{,}883$). **(A)** Hexbin plot of predicted versus observed editing scores, with the dashed diagonal indicating identity and the red line showing the linear fit (Spearman $\rho = 0.8145$, Pearson $r = 0.7922$). **(B)** Residuals plotted against predicted editing scores; the black line marks zero residual and the red line shows a rolling average trend (mean residual $= -0.8349$, residual standard deviation $= 3.5561$). **(C)** Overlaid density distributions of observed and predicted editing scores, showing the low-efficiency enrichment present in the DeepPrime test set. **(D)** Absolute-error distribution on a logarithmic frequency axis (RMSE = 3.653, mean absolute error = 1.971, median absolute error = 0.643).

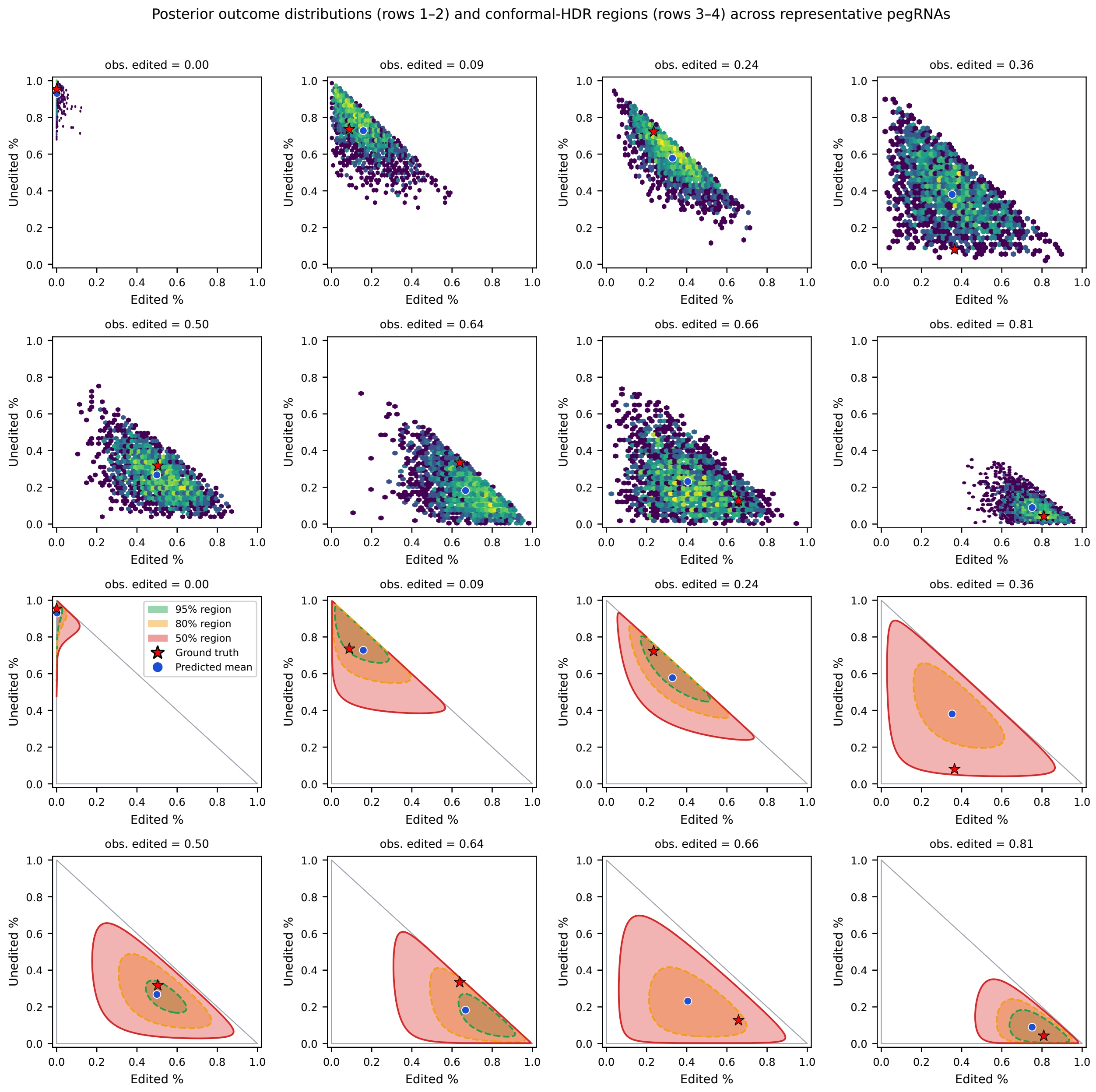


Figure S2: Example posterior outcome distributions and conformal-HDR regions for representative pegRNA predictions. **Top two rows:** Posterior samples drawn from the Dirichlet distribution projected onto the 2D outcome simplex (edited fraction vs. unedited fraction); colour intensity reflects density. **Bottom two rows:** Corresponding 50% (red), 80% (orange), and 95% (green) split-conformal HDR regions, each calibrated on the held-out validation fold ($n_{\mathrm{cal}} = 13{,}828$). Regions are super-level sets of $-\log \mathrm{Dir}(y; \hat{\boldsymbol{\alpha}})$ on the simplex and remain inside the triangular simplex projection by construction. Ground truth (red star) and predicted mean (blue circle) overlaid. Panels span low, intermediate, and high observed editing efficiency.

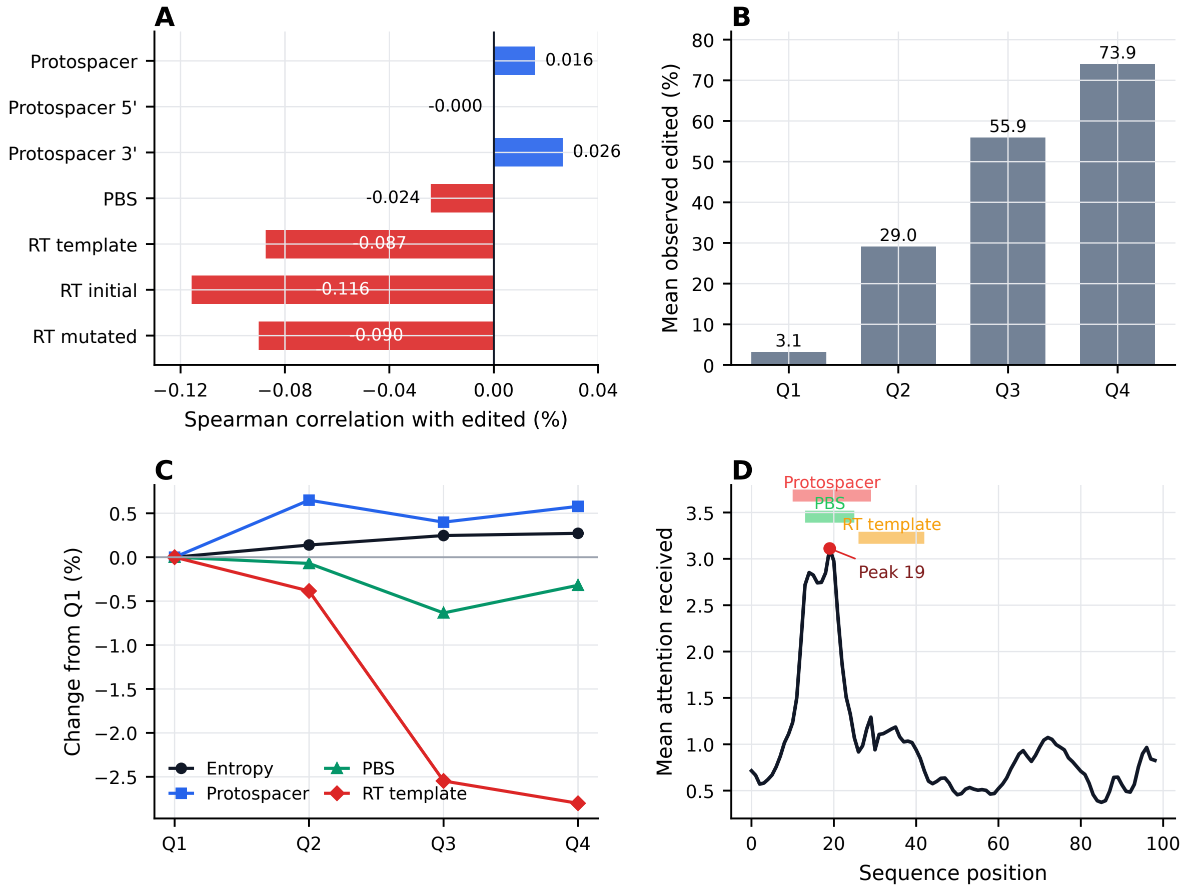


Figure S3: Attention-based diagnostics for crispAIPE on the target-disjoint PRIDICT Library-1 test set ($n = 13{,}844$). **(A)** Spearman correlations between regional attention summaries and observed edited efficiency: protospacer attention is near zero overall (all protospacer positions $\rho = 0.016$; positions 10–19 $\rho \approx 0.000$; positions 20–29 $\rho = 0.026$), PBS attention is slightly negative ($\rho = -0.024$), and RT-template attention is negative across the full template ($\rho = -0.087$), RT-initial segment ($\rho = -0.116$), and RT-mutated segment ($\rho = -0.090$). **(B)** Mean observed editing efficiency across quartiles used for attention summaries. **(C)** Relative changes in attention entropy and regional attention summaries across efficiency quartiles, normalized to Q1. **(D)** Position-wise average attention, with shaded bands marking protospacer, PBS, and RT-template intervals.

#### Supplementary Tables

Table S1: Architecture ablation on the mutation-level target-disjoint PRIDICT Library-1 split. All models were retrained with the same train/validation/test partition and Dirichlet objective. The hybrid transformer–CNN model outperforms both single-branch variants across edited, unedited, indel, and overall rank correlations.

| Architecture | Val loss | Test loss | Edited $\rho$ | Unedited $\rho$ | Indel $\rho$ | Overall $\rho$ |
|---|---|---|---|---|---|---|
| Transformer-only | -2.3027 | -2.2432 | 0.6204 | 0.6825 | 0.5499 | 0.6964 |
| CNN-only | -2.6213 | -2.6423 | 0.7884 | 0.8049 | 0.6320 | 0.8146 |
| Hybrid transformer–CNN | **-2.8517** | **-2.8385** | **0.8352** | **0.8433** | **0.6930** | **0.8507** |

Table S2: Outcome-parameterisation ablation on the mutation-level target-disjoint PRIDICT Library-1 split. Rank correlations are comparable across rows; validation and test objectives are not shown because the Dirichlet negative log-likelihood, softmax cross-entropy, and logit-normal approximation optimise different losses. Softmax gives marginally higher point-prediction rank correlation but does not provide the Dirichlet posterior concentration parameters used for uncertainty diagnostics.

| Parameterisation | Objective | Edited $\rho$ | Unedited $\rho$ | Indel $\rho$ | Overall $\rho$ |
|---|---|---|---|---|---|
| Dirichlet concentration | Dirichlet NLL | 0.8352 | 0.8433 | 0.6930 | 0.8507 |
| Softmax point head | Cross-entropy | **0.8425** | **0.8496** | **0.6975** | **0.8563** |
| Logit-normal head | Logit-space approximation | 0.8201 | 0.8237 | 0.5974 | 0.8290 |

Table S3: PRIDICT Library-2 zero-shot transfer and small-sample cell-line adaptation. Values are mean Spearman correlations across five repeated held-out splits. Test-set sizes vary slightly by split because target-level grouping was preserved. Head fine-tuning used the Library-1-trained crispAIPE checkpoint with limited target-cell training data.

| Condition | Mean held-out $n$ | Zero-shot $\rho$ | Head fine-tuned $\rho$ |
|---|---|---|---|
| HEK293T PE2 Library-2 | 277.0 | 0.766 | 0.895 |
| U2OS PE2 Library-2 | 264.8 | 0.340 | 0.837 |
| K562 PE2 Library-2 | 253.8 | 0.456 | 0.860 |
| Liver GFP-positive PE2-Adeno Library-2 | 290.0 | 0.384 | 0.785 |

#### Supplementary Note 1: Split-Conformal HDR Construction

We construct prediction regions for the compositional outcome $\mathbf{y} \in \Delta^2$ using split conformal prediction [11, 7, 2] with the Dirichlet highest-density region (HDR) geometry of Hyndman [6], following Amaral et al. [1]. The construction has two parts: a region geometry (Dirichlet level set) and a calibration step (one global threshold from a held-out fold).

**Algorithm 1: Split-Conformal HDR**

**Input:** Calibration fold $\{(x_i, y_i^{\text{obs}})\}_{i=1}^{n_{\text{cal}}}$, trained model $\hat{\boldsymbol{\alpha}}(\cdot)$, nominal level $\gamma \in (0,1)$, test input $x^*$.
**Output:** Prediction region $C(x^*) \subset \Delta^2$ with $\mathbb{P}(y^{\text{obs}} \in C(X^*)) \geq 1-\gamma$ under exchangeability.

**// Calibration phase (run once)**
**for** $i = 1$ to $n_{\text{cal}}$ **do**
  $s_i \leftarrow -\log \mathrm{Dir}(y_i^{\text{obs}}; \hat{\boldsymbol{\alpha}}(x_i))$ *// Nonconformity score*
**end for**
$\hat{q} \leftarrow$ empirical $\lceil (n_{\text{cal}}+1)(1-\gamma) \rceil / n_{\text{cal}}$ quantile of $\{s_i\}_{i=1}^{n_{\text{cal}}}$

**// Inference for a test input $x^*$**
Compute $\hat{\boldsymbol{\alpha}}^* = \hat{\boldsymbol{\alpha}}(x^*)$
$C(x^*) \leftarrow \{y \in \Delta^2 : -\log \mathrm{Dir}(y; \hat{\boldsymbol{\alpha}}^*) \leq \hat{q}\}$
**return** $C(x^*)$

**Region geometry.** For fixed $\hat{\boldsymbol{\alpha}}$, the region $C(x^*)$ is a super-level set of the Dirichlet density: it contains the highest-density points under $\mathrm{Dir}(\hat{\boldsymbol{\alpha}}^*)$. By construction it stays inside the simplex $\Delta^2$ and is the smallest-volume Bayesian credible region at any given mass under $\mathrm{Dir}(\hat{\boldsymbol{\alpha}}^*)$ [6]. The threshold $\hat{q}$ shifts the level set inward or outward according to held-out validation evidence rather than the model's own posterior.

**Coverage guarantee.** Split conformal prediction provides finite-sample distribution-free marginal coverage:

$$\mathbb{P}\big(\, y_{\text{test}}^{\text{obs}} \in C(x_{\text{test}}) \,\big) \geq 1-\gamma, \tag{1}$$

under exchangeability of $\{(x_i, y_i^{\text{obs}})\}_{i=1}^{n_{\text{cal}}} \cup \{(x_{\text{test}}, y_{\text{test}}^{\text{obs}})\}$ [11, 7]. The guarantee holds for any choice of nonconformity score; here the score is the negative Dirichlet log-density at the observed label, which means a well-calibrated Dirichlet head yields the smallest regions, while a miscalibrated head still satisfies the coverage guarantee but pays for it in region width.

**Calibration fold.** We use the mutation-disjoint validation fold of PRIDICT Library-1 ($n_{\text{cal}} =$ 13,828 pegRNAs) for calibration, which is disjoint from the training fold used to fit $\hat{\boldsymbol{\alpha}}(\cdot)$ and from the test fold used for evaluation ($n_{\text{test}} =$ 13,844). The empirical quantile $\hat{q}$ at $\gamma = 0.95$ on the canonical checkpoint is $\hat{q}_{0.95} \approx 1.15$.

**Region area as a reliability indicator.** The 2D-projected area of $C(x^*)$ on the (edited, unedited) simplex projection is estimated by Monte Carlo: we draw $M =$ 8,000 samples uniformly from $\Delta^2$ (i.e., from $\mathrm{Dir}(1,1,1)$), evaluate $-\log \mathrm{Dir}(y_j; \hat{\boldsymbol{\alpha}}^*)$ at each, and take the empirical inside-fraction times the area of the triangular projection (0.5):

$$\widehat{A}(\hat{\boldsymbol{\alpha}}^*, \hat{q}) = \frac{0.5}{M} \sum_{j=1}^{M} \mathbf{1}[\, -\log \mathrm{Dir}(y_j; \hat{\boldsymbol{\alpha}}^*) \leq \hat{q}\,]\,. \tag{2}$$

$\widehat{A}$ is a per-pegRNA scalar that summarises predicted uncertainty and, on the canonical test fold, has Spearman correlation $\rho = 0.43$ with absolute prediction error (main-text Fig. 3C). Lower $\widehat{A}$ indicates a sharper region and, empirically, lower expected error.

**Region construction ablation.** We additionally evaluated two simpler constructions on the same target-disjoint test fold (Supplementary Fig. S4, Supplementary Table S4): (i) the Bayesian HDR of Hyndman, which replaces $\hat{q}$ with the per-test-input $(1-\gamma)$ sample quantile of $\log \mathrm{Dir}(y; \hat{\boldsymbol{\alpha}}^*)$ over $S = 1000$ posterior samples, and (ii) the PCA ellipse baseline used in earlier work [10], which approximates the region as a 2D Gaussian ellipse with semi-axes $r_i = \sqrt{\chi^2_{2,\gamma} \lambda_i}$ where $\lambda_i$ are the eigenvalues of the closed-form 2D Dirichlet marginal covariance. Both alternatives provide Bayesian credible regions only and inherit the marginal miscalibration of the Dirichlet head. The conformal HDR is the only construction that empirically matches the nominal joint coverage level on the test fold, motivating its adoption as the headline construction.

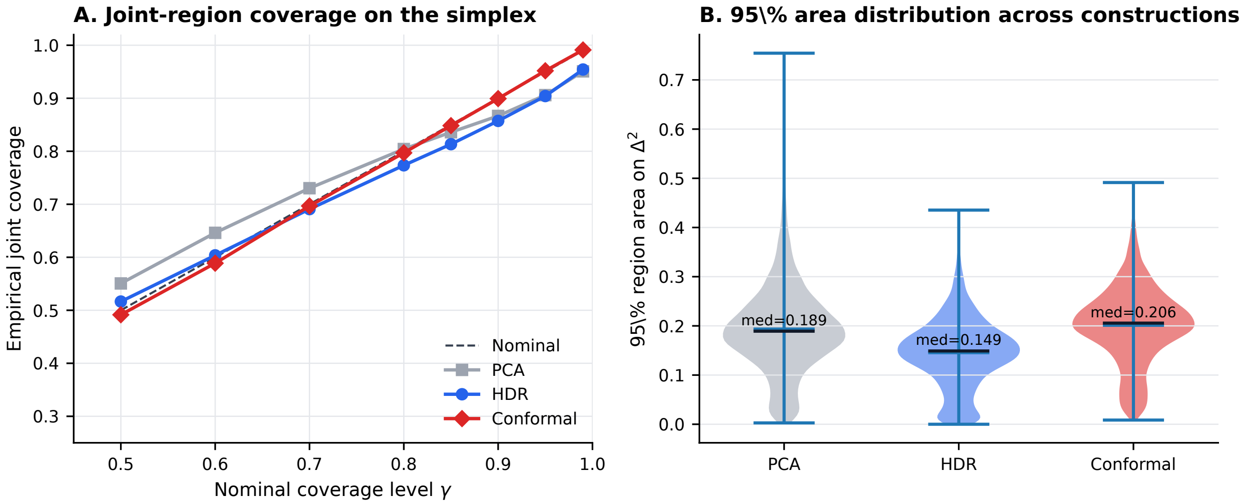


Figure S4: Region-construction ablation on the target-disjoint PRIDICT Library-1 test set ($n =$ 13,844), with calibration on the held-out validation fold ($n_{\text{cal}} =$ 13,828). **(A)** Empirical joint coverage on the simplex as a function of nominal level $\gamma$ for three constructions: the PCA ellipse baseline (grey), the Bayesian HDR of Hyndman (blue), and the split-conformal HDR (red). The dashed black line is nominal $y = \gamma$. The conformal HDR matches the nominal level across $\gamma \in \{0.50, 0.60, \ldots, 0.99\}$; PCA and Bayesian HDR drift below nominal at $\gamma = 0.95$ and above nominal in mid-range, reflecting the marginal miscalibration of the Dirichlet head. **(B)** Distribution of 95% region areas on the (edited, unedited) projection of the simplex (maximum possible area 0.5). Bayesian HDR is the sharpest at fixed nominal level (median 0.149) but inherits the model's miscalibration; conformal HDR is wider (median 0.206) because it pays for calibration; the PCA ellipse (median 0.189) sits between but can extend outside the simplex.

Table S4: Region-construction ablation on the target-disjoint PRIDICT Library-1 test set. Empirical joint coverage and median 95% region area on the (edited, unedited) projection ($n = 13{,}844$ test pegRNAs; conformal HDR calibrated on $n_{\text{cal}} = 13{,}828$ validation pegRNAs). The third column reports Spearman correlation between per-pegRNA region area and absolute prediction error at $\gamma = 0.95$, an empirical check that the region area carries reliability information.

| Construction | $\gamma$ | Joint coverage | Median area | Spearman $\rho$(area, \|err\|) |
|---|---|---|---|---|
| PCA ellipse | 0.95 | 0.906 | 0.189 | 0.445 |
| Bayesian HDR (Hyndman) | 0.95 | 0.904 | 0.149 | 0.428 |
| **Conformal HDR (Amaral)** | 0.95 | **0.952** | 0.206 | 0.431 |
| PCA ellipse | 0.90 | 0.867 | 0.145 | 0.445 |
| Bayesian HDR (Hyndman) | 0.90 | 0.857 | 0.118 | 0.423 |
| **Conformal HDR (Amaral)** | 0.90 | **0.899** | 0.148 | 0.416 |
| PCA ellipse | 0.80 | 0.804 | 0.102 | 0.445 |
| Bayesian HDR (Hyndman) | 0.80 | 0.773 | 0.086 | 0.416 |
| **Conformal HDR (Amaral)** | 0.80 | **0.797** | 0.097 | 0.387 |
| PCA ellipse | 0.50 | 0.551 | 0.044 | 0.445 |
| Bayesian HDR (Hyndman) | 0.50 | 0.516 | 0.038 | 0.404 |
| **Conformal HDR (Amaral)** | 0.50 | **0.492** | 0.031 | −0.060 |

#### Supplementary Note 2: Encoding and Tokenization Details

We employ a hybrid encoding strategy combining sequence-based features with positional and functional annotations to capture the multi-faceted nature of prime editing. For nucleotide encoding, we use 5-symbol nucleotide/gap encoding representing the alphabet $\{A, T, G, C, \text{gap}\}$, where gaps accommodate insertions and deletions. Wild-type and mutated sequences are represented across a fixed sequence length $L$.

To capture editing-specific information, we merge the wild-type and mutated sequence matrices through element-wise OR operations, yielding a $5 \times L$ matrix that highlights positions where edits occur. However, OR operations alone cannot preserve edit directionality (e.g., distinguishing $A \rightarrow G$ from $G \rightarrow A$ substitutions). To address this, we concatenate a 2-bit direction channel encoding the transformation type: '00' for matches, '01' for wild-type-to-mutant transitions, '10' for mutant-to-wild-type transitions, and '11' for insertion/deletion events, yielding a $7 \times L$ representation capturing both edit positions and their directionality.

We further augment this representation with four binary location channels explicitly marking functionally critical pegRNA regions: (i) protospacer, (ii) PBS, (iii) RTT-initial, and (iv) RTT-mutated positions. Each channel is a $1 \times L$ binary vector with '1' at relevant positions and '0' elsewhere. The final unified representation combines all components into an $11 \times L$ matrix ($d = 5 + 2 + 4 = 11$). The wild-type nucleotide sequence is separately converted to token indices and processed through a learnable embedding layer with $d_{\text{embed}} = 64$ before concatenation with the unified representation. No k-mer tokenization is used in the reported crispAIPE model.

#### Supplementary Note 3: Glossary of Specialised Terms

Definitions provided for non-specialist readers; primary references in Section 7 of the bibliography.

Table S5: Glossary of prime-editing terms used in the main text. Full chemistry and citations are deferred to the primary literature [3, 4, 9, 5, 12, 8].

| Term | Definition |
|---|---|
| Cas9 nickase (nCas9, H840A) | A Cas9 variant in which the HNH nuclease is disabled by the H840A mutation, so the editor cuts only one DNA strand. Prime editors use the nickase form to avoid double-strand breaks. |
| PAM | Protospacer adjacent motif (NGG for *S. pyogenes* Cas9); a short DNA motif immediately $3'$ of the protospacer on the non-target strand. Required for Cas9 binding; the PAM lives in the genome, not the guide. |
| Protospacer | The $\sim 20$ bp stretch of genomic DNA, $5'$ of a PAM, that a guide RNA targets. |
| Spacer | The $\sim 20$ nt portion of the guide RNA whose sequence matches the protospacer (with T→U) and base-pairs with the target strand of the DNA. |
| sgRNA | Single-guide RNA: an engineered $\sim 100$ nt RNA combining bacterial crRNA and tracrRNA; $5'$ $\sim 20$ nt spacer plus $\sim 80$ nt invariant scaffold that engages Cas9. |
| pegRNA | Prime editing guide RNA: an extended sgRNA whose $3'$ end carries an additional RTT and PBS that together encode the edit. |
| PBS | Primer binding site, the $3'$ pegRNA segment ($\sim$ 8–17 nt) that hybridises to the nicked PAM-containing strand and provides the $3'$-OH primer for reverse transcription. |
| RTT | Reverse transcription template ($\sim$ 10–20 nt) that templates the new DNA flap, encoding the intended edit plus homology to the unedited $3'$ genome. |
| RTT overhang | Portion of the RTT extending $3'$ of the intended edit; provides homology that allows the new flap to invade and replace the original sequence. Independent design variable that this work treats explicitly via target-disjoint splitting. |

Table S5 (continued)

| Term | Definition |
|---|---|
| $3'$ / $5'$ flap | Single-stranded DNA branches created at the nick after reverse transcription. The $3'$ flap carries the new sequence; the $5'$ flap is the displaced original. FEN1-style endonucleases preferentially cleave the $5'$ flap; ligation installs the $3'$ flap. |
| ngRNA | Nicking guide RNA used in PE3/PE3b to direct a second nick to the unedited strand and bias mismatch repair toward the edited strand. |
| epegRNA | Engineered pegRNA carrying a $3'$ structured motif (e.g., tevopreQ1, mpknot) that protects the pegRNA from $3' \to 5'$ exonucleases; typically 3–4-fold efficiency gain. |
| MMR | Mismatch repair pathway (MutS$\alpha$, MutL$\alpha$). In prime editing, MMR preferentially removes the edited strand and reverts the edit; PE4/PE5 use a dominant-negative MLH1 to transiently inhibit MMR. |
| PE1 / PE2 / PE3 / PE3b | Anzalone 2019: PE1 = Cas9 nickase + wild-type M-MLV RT; PE2 = pentamutant M-MLV RT (more efficient); PE3 = PE2 + ngRNA on the unedited strand; PE3b = PE3 where the ngRNA only nicks after the edit is installed. |
| PE4 / PE5 / PEmax | Chen 2021: PE2 + MLH1dn (PE4) and PE3 + MLH1dn (PE5). PEmax is the optimised PE2 protein backbone used in PE4/PE5/PE6/PE7. |
| PE6 (a–g) | Doman 2023: phage-evolved compact RT variants (PE6a–d) and PEmax-sized editors with engineered Cas9 mutations (PE6e–g); enable AAV delivery. |
| PE7 | Yan 2024: PEmax fused to the N-terminal La motif of human La/SSB protein, which protects pegRNA $3'$ ends; large gains in primary cells. |
| Dirichlet distribution | A multivariate distribution over the simplex; the natural likelihood for vectors of proportions that sum to one. Here, the 3-D Dirichlet over (edited, unedited, indel) captures both the predicted mean and predicted uncertainty per pegRNA. |
| Conformal prediction (split-conformal HDR) | A distribution-free framework that calibrates prediction regions to a user-chosen coverage level using a held-out calibration fold, with finite-sample coverage under exchangeability. Used here to construct calibrated highest-density regions on the editing-outcome simplex. |

### Supplementary Figures: Architecture and Cell-context Analyses

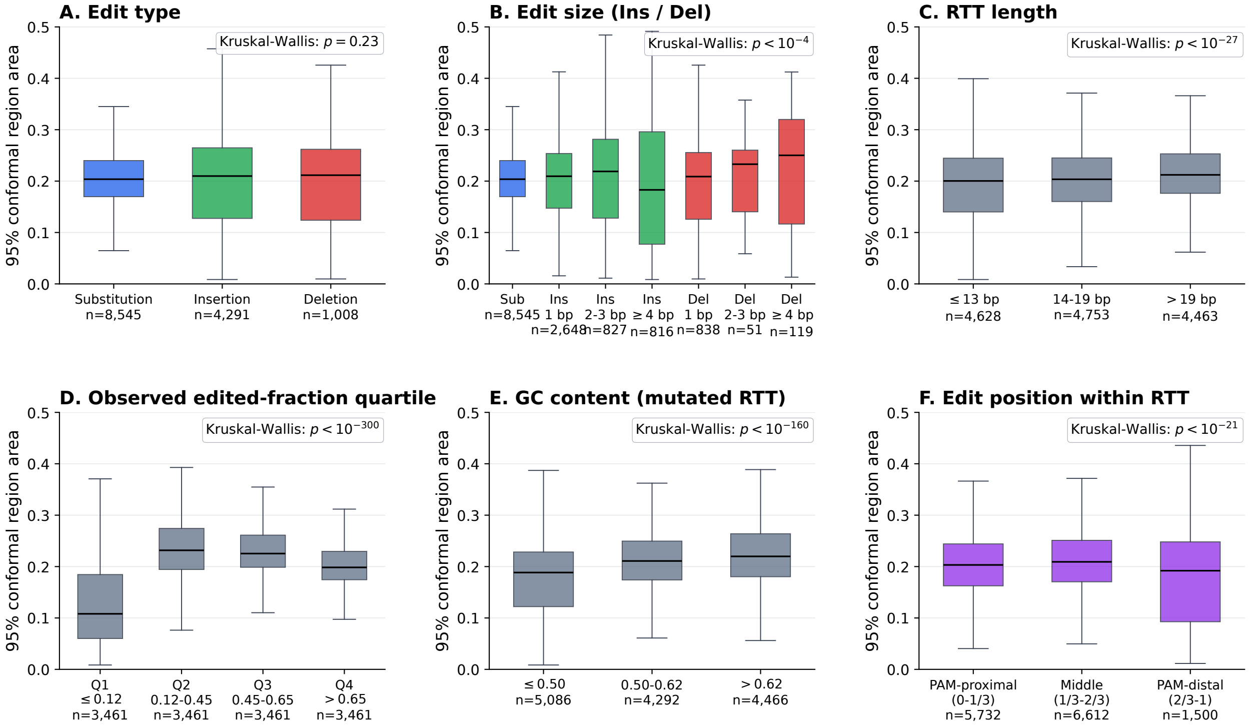


Figure S5: pegRNA architectural and edit-context determinants of prediction uncertainty on the target-disjoint Library-1 test set ($n = 13{,}844$). Each panel stratifies the per pegRNA–target pair 95% split-conformal HDR region area on the (edited, unedited) simplex projection by a different biological factor; box plots show median, interquartile range, and 1.5 IQR whiskers (outliers suppressed). Per-stratum sample sizes are reported under each tick label and Kruskal–Wallis p-values are annotated in each panel. **(A)** Edit type (substitution / insertion / deletion). **(B)** Edit size, with insertions and deletions sub-stratified by length bins (1, 2–3, $\geq 4$ bp); substitutions shown for reference at size 1. **(C)** RTT length tertile. **(D)** Observed edited-fraction quartile (PBS length is uniformly 12 bp in PRIDICT Library-1 and is therefore not stratifiable); used to test whether the model is most uncertain at intermediate editing efficiencies. **(E)** GC content of the mutated RTT, in tertiles. **(F)** Edit position within the RTT, terciled by the normalised first-divergence position between WT and mutated RTT (PAM-proximal = closer to the priming site, PAM-distal = far end).

Calibration guarantees address the population-level coverage question but say nothing about which pegRNA architectures the model is more or less certain about. We stratified the per pegRNA–target pair 95% conformal-HDR region area by six biological factors on the target-disjoint test set (Fig. S5). Edit type alone (Sub / Ins / Del) did not separate uncertainty (Kruskal–Wallis $p = 0.23$; Fig. S5A), but edit size did: large deletions ($\geq 4$ bp) carried the largest median region area (0.25 versus 0.20 for single-base substitutions; $p \approx 7 \times 10^{-5}$; Fig. S5B). Longer RTTs were associated with larger regions (median area 0.212 for the longest tertile versus 0.200 for the shortest; $p \approx 3 \times 10^{-28}$; Fig. S5C). Region area also varied strongly with the observed edited-fraction quartile (Fig. S5D): pegRNAs in the lowest observed-edited quartile (Q1, edited fraction $\leq 0.04$) had the lowest median region area (0.11), consistent with the model confidently predicting that no editing will occur;

uncertainty peaked in the central quartiles (Q2–Q3, $\approx 0.23$) where the outcome is genuinely most ambiguous, and partially recovered for high-efficiency pegRNAs. GC content of the mutated RTT was the dominant biological correlate of uncertainty (Fig. S5E): low-GC RTTs gave a median region area of 0.188 versus 0.220 for high-GC RTTs ($p \approx 4 \times 10^{-161}$). Finally, the position of the edit within the RTT (Fig. S5F) carried a smaller but highly significant effect ($p \approx 5 \times 10^{-22}$), with PAM-proximal and mid-RTT edits showing larger region areas than PAM-distal edits. Together, these stratifications turn the global calibration claim into a feature-level uncertainty map: GC content, edit size for deletions, and RTT length are the architectural variables that most reliably forecast where crispAIPE expects per pegRNA–target pair outcomes to be more uncertain.

Prime editing efficiency depends on repair context, expression system, and cellular state. We evaluated the Library-1-trained model on the four PRIDICT Library-2 cell-context datasets and performed repeated small-sample target-cell adaptation using head fine-tuning (Supplementary Table S3). Zero-shot transfer was strongest for matched HEK293T PE2 data (mean Spearman $\rho = 0.766$ across five held-out splits) and weaker for U2OS, K562, and liver GFP-positive PE2-Adeno contexts (mean $\rho = 0.340$–$0.456$); limited target-cell fine-tuning improved performance substantially across all evaluated contexts (mean $\rho = 0.785$–$0.895$).

These zero-shot averages, however, mask a strongly heterogeneous within-dataset structure that the calibrated conformal regions expose. Reusing the Library-1 validation-fold quantile $\hat{q}_{0.95} = 1.15$ unchanged, we computed the per pegRNA–target pair 95% region area on each Library-2 dataset and asked whether filtering pegRNAs by region area improves zero-shot accuracy in each cell line (Fig. S6). For every cell line, retaining the top-confident fraction by ascending region area produced a monotonic improvement in Spearman $\rho$ on the edited fraction (Fig. S6A): from 0.667 to 0.883 in HEK293T PE2 as retention decreased from 100% to 25%; from 0.495 to 0.752 in U2OS PE2; from 0.484 to 0.731 in K562 PE2; and from 0.352 to 0.581 in liver GFP-positive PE2-Adeno. Two reference filters confirm that this gain is specific to the conformal-region signal (Fig. S6A): ranking by the Dirichlet concentration $\alpha_0$ alone produces essentially flat curves (e.g., $0.495 \to 0.533$ in U2OS at 25% retention), and a random subsample baseline tracks the full-set Spearman to within $\pm 0.01$ at every retention level in every cell line. The same comparison summarised at the 25% retention operating point (Fig. S6B) shows region-area filtering delivers $\Delta\rho \approx +0.22$ to $+0.26$ across all four cell contexts, while $\alpha_0$ and random filters cluster within $\pm 0.06$ of zero. Generalising the small-$N$ illustration in Fig. S6D, the top-$N$ versus bottom-$N$ MAE curves across $N \in [5, 100]$ (Fig. S6E) show a wide and persistent gap between the most- and least-confident designs in every cell line, with the gap narrowing as $N$ grows toward the full set. Per-sample, the conformal region area was most strongly associated with absolute zero-shot error in the matched HEK293T context (Spearman $\rho = 0.43$; Fig. S6C). Comparing the most- and least-confident designs for U2OS PE2 (Fig. S6D), the top-5 pegRNAs ranked by Library-1 region area show substantially lower observed error than the bottom-5. crispAIPE's calibrated uncertainty therefore functions as an actionable filter for cross-context pegRNA design: the model is not cell-line invariant, but its Library-1-calibrated region area selects the subset of designs whose Library-1 prediction is most likely to carry over to a new cell context.

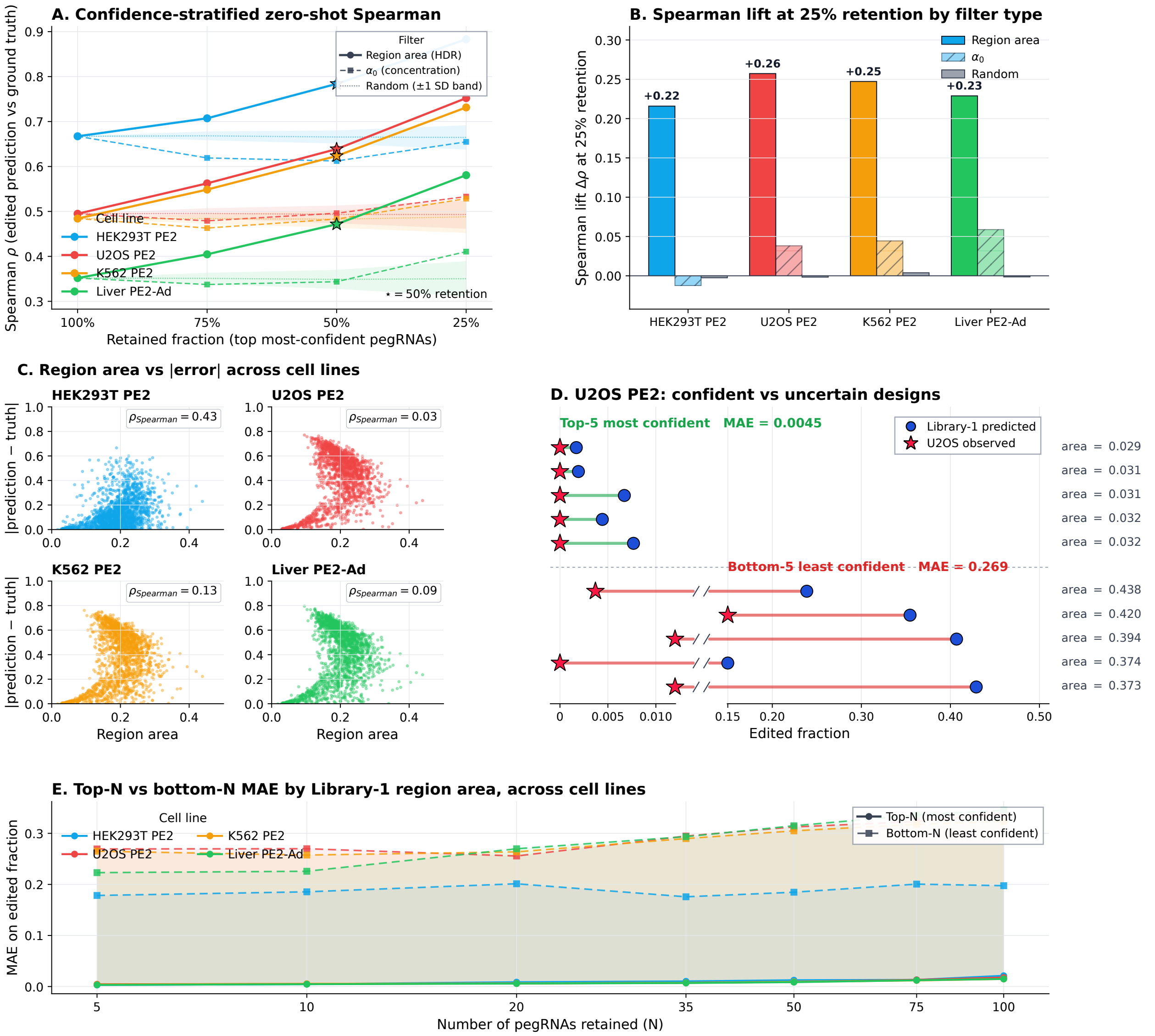


Figure S6: Library-1-calibrated conformal region area predicts which pegRNAs transfer well across PRIDICT Library-2 cell contexts. The calibration quantile $\hat{q}_{0.95} = 1.15$ from the Library-1 validation fold is reused unchanged. **(A)** Confidence-stratified zero-shot Spearman $\rho$ between predicted and observed edited fraction in each cell line, computed after retaining the top $\{100, 75, 50, 25\}\%$ of pegRNAs by ascending region area (most-confident first; solid lines). Two reference filters are overlaid for comparison: ranking by the Dirichlet concentration $\alpha_0$ (dashed lines; scalar model-internal confidence) and uniform random subsampling (dotted lines; shaded band shows mean $\pm$ 1 SD over $B = 200$ replicates). Stars mark the 50% retention operating point. **(B)** Spearman lift $\Delta\rho$ at 25% retention versus the full set, decomposed by filter type: region area (solid colored bars), $\alpha_0$ (hatched bars), and random subsampling (grey bars). Region-area filtering yields $\Delta\rho \approx +0.22$ to $+0.26$ across all four cell contexts; the two reference filters cluster near zero. **(C)** Conformal region area versus absolute zero-shot prediction error per cell line; per-panel Spearman correlation annotated. **(D)** U2OS PE2 confident versus uncertain designs: top-5 and bottom-5 pegRNAs sorted by Library-1-calibrated region area, with the corresponding Library-1 predicted edited fraction and U2OS observed edited fraction. The top-5 confident pegRNAs show substantially lower MAE in U2OS than the bottom-5. **(E)** Quantitative top-$N$ versus bottom-$N$ MAE on edited fraction by Library-1 region area, for $N \in \{5, 10, 20, 35, 50, 75, 100\}$, across all four cell contexts. Solid lines are the most-confident $N$ pegRNAs, dashed lines are the least-confident $N$; the shaded gap between curves at each $N$ measures the operational benefit of region-area filtering.
